# Retrieval Augmented Protein Language Models for Protein Structure Prediction

**DOI:** 10.1101/2024.12.02.626519

**Authors:** Pan Li, Xingyi Cheng, Le Song, Eric Xing

**Author notes:** Equal contribution (Pan Li, Xingyi Cheng). Correspondence to: Le Song < >, Eric Xing < >. Proceedings of the 42 ^nd^ International Conference on Machine Learning*, Vancouver, Canada. PMLR 267, 2025.

## Abstract

The advent of advanced artificial intelligence technology has significantly accelerated progress in protein structure prediction, with AlphaFold2 setting a new benchmark for prediction accuracy by leveraging the Evoformer module to automatically extract co-evolutionary information from multiple sequence alignments (MSA). To address AlphaFold2’s dependence on MSA depth and quality, we propose two novel models: AIDO.RAGPLM and AIDO.RAGFold, pretrained modules for **R**etrieval-**A**u**G**mented protein language model and structure prediction in an AI-driven Digital Organism (Song et al., 2024). AIDO.RAGPLM integrates pre-trained protein language models with retrieved MSA, surpassing single-sequence protein language models in perplexity, contact prediction, and fitness prediction. When sufficient MSA is available, AIDO.RAGFold achieves TM-scores comparable to AlphaFold2 while operating up to eight times faster, and significantly outperforms AlphaFold2 when MSA is insufficient (ΔTM-score=0.379, 0.116 and 0.059 for 0, 5 and 10 MSA sequences as input). Additionally, we developed an MSA retriever using hierarchical ID generation that is 45 to 90 times faster than traditional methods, expanding the MSA training set for AIDO.RAGPLM by 32%. Our findings suggest that AIDO.RAGPLM provides an efficient and accurate solution for protein structure prediction, particularly in scenarios with limited MSA data. The AIDO.RAGPLM model has been open-sourced and is available on https://huggingface.co/genbio-ai/AIDO.Protein-RAG-3B.

## 1. Introduction

The advent of advanced artificial intelligence technology has significantly accelerated progress in protein structure prediction. AlphaFold2 (Jumper et al., 2021), a pioneering method in this field, has set a new benchmark for prediction accuracy. Multiple sequence alignment (MSA) plays a crucial role in protein structure prediction. Unlike previous methods that required manual calculation of MSA features (Senior et al., 2020), AlphaFold2 leverages the Evoformer module to automatically extract co-evolutionary information from MSA, thereby enhancing the efficiency of information utilization.

However, the efficacy of structure prediction methods like AlphaFold2 is heavily dependent on the depth and quality of the MSA. Consequently, it is imperative to prepare an extensive sequence database. When the number of homologous sequences is insufficient, the performance of AlphaFold2 deteriorates significantly. To address this limitation, methods based on large-scale pre-trained protein language models have been proposed. For instance, ESMFold (Lin et al., 2023), OmegaFold (Wu et al., 2022), ESM3 (Hayes et al., 2024) and xTrimoPGLM-Fold (Chen et al., 2024b) have demonstrated commendable results using a single sequence as input. Nevertheless, even with 100-billion parameters, models like xTrimoPGLM and ESM3 remain inferior to AlphaFold2 in structure prediction when MSA is used as input, underscoring the importance of MSA. Although several PLM have attempted to integrate multiple sequences for training (see Appendix A), there is currently no validation for using retrieved augmented PLM for end-to-end protein structure prediction.

In this paper, we integrate pre-trained protein language models with retrieved MSA to propose a novel approach termed Protein Language Model with Retrieved Augmented MSA (RAGPLM) (see Figure 1). This approach allows for the incorporation of co-evolutionary information from MSA in structure prediction while compensating for insufficient MSA information through large-scale pre-training. We concatenate the query sequence with aligned homologous sequences into a long sequence (up to 12.8k) and perform pre-training by column span mask strategy based on a transformer encoder framework. Our method surpasses single-sequence protein language models in perplexity, contact prediction, and fitness prediction. Subsequently, we utilized AIDO.RAGPLM as a feature extractor, integrating it with the folding trunks and Structure Modules to achieve end-to-end structural prediction (AIDO.RAGFold). Our findings indicate that when sufficient MSA is available, our method achieves results comparable to AlphaFold2 and is eight times faster; when MSA is insufficient, our method significantly outperforms AlphaFold2.

**Figure 1.**
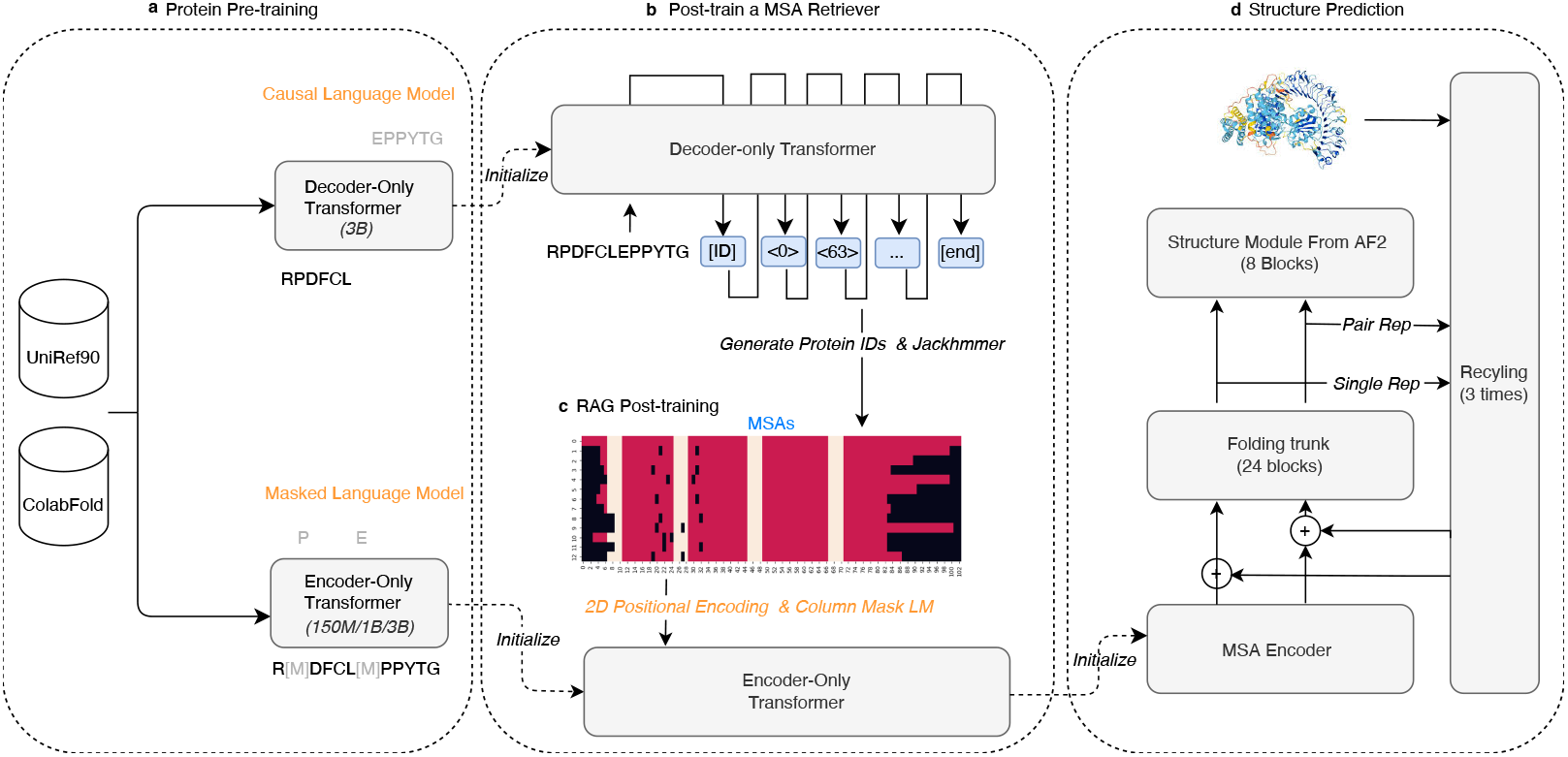
Schematic diagram of MSA retriever, AIDO.RAGPLM, and AIDO.RAGFold. (a) A decoder-only and an encoder-only transformer model are trained using UniRef90 and ColabFold protein sequence databases with CLM and MLM losses, respectively. (b) The MSA Retriever is fine-tuned on an MSA dataset to generate MSA sequence IDs from a query sequence, enabling the creation of a UniRef50 MSA training dataset. (c) AIDO.RAGPLM is trained on the UniRef50 MSA dataset using column span masking and recovery loss. (d) AIDO.RAGPLM acts as a feature extractor for protein structure prediction.

To expedite MSA acquisition, we also developed an MSA retriever using hierarchical ID generation. This retriever is 45 to 90 times faster than traditional HHblits (Steinegger et al., 2019) in MSA retrieval, which is used to expand the MSA training set for AIDO.RAGPLM by 32%.

## 2. Methods

Our method consists of three major components, MSA retriever, AIDO.RAGPLM and AIDO.RAGFold, which we explain more details below.

### 2.1. MSA retriever

Searching for multiple sequence alignments (MSAs) in large sequence databases is time-consuming. Inspired by (Wang et al., 2023) that generates relevant document identifiers by sequence-to-sequence network in document retrieval, we developed an MSA retriever to generate hierarchical identifiers for homologous sequences for a query protein sequence (see Figure 3). The protocol comprises three steps: (1) Construct hierarchical IDs for each sequence in UniClust30 (UC30) (Mirdita et al., 2016) through hierarchical K-means clustering of embedding; (2) Fine-tune a pretrained casual language model with 3-billion parameters (CLM-3B, (Cheng et al., 2024)) to memorize the ID of each sequence on UC30 dataset; (3) Continue to fine-tune the model to generalize to IDs of homologous sequences on the HHblits MSA dataset. For detailed training information, please refer to Appendix C. During inference, the MSA retriever generates each ID token sequentially, which corresponds to the nodes of the tree, until the UC30 node is reached. We perform multiple generations using different parameters and aggregate all retrieved sequences. Jackhmmer (Johnson et al., 2010) is then used to filter and align the homologous sequences. We use MSA Retriever to expand the MSA training data for AIDO.RAGPLM (see Appendix D).

**Figure 2.**
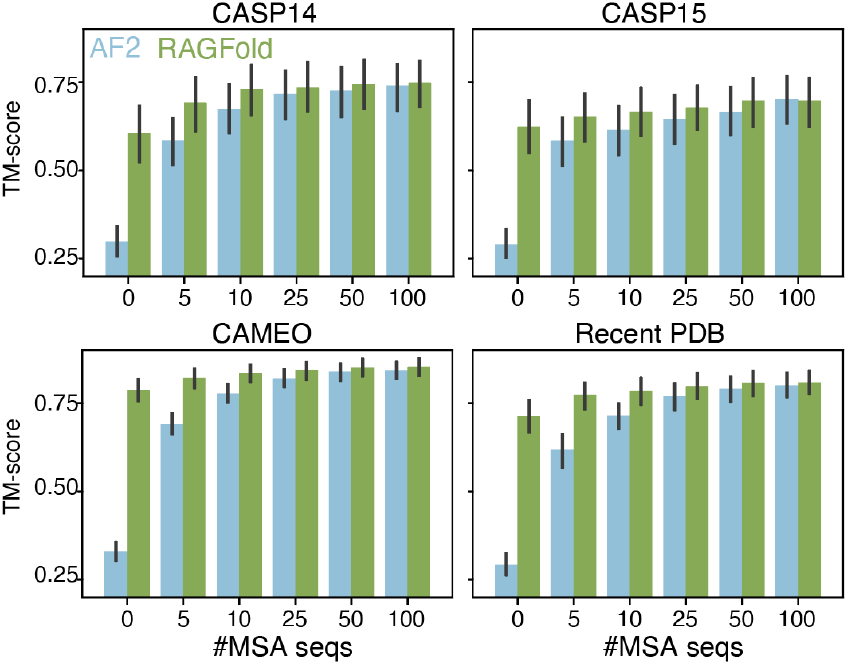
TM-scores of AlphaFold2 and AIDO.RAGFold on four test datasets with limited MSA sequences as input. AlphaFold2 and AIDO.RAGFold are represented by blue and green bars respectively. The *x* axis represents the upper bound of the MSA number.

**Figure 3.**
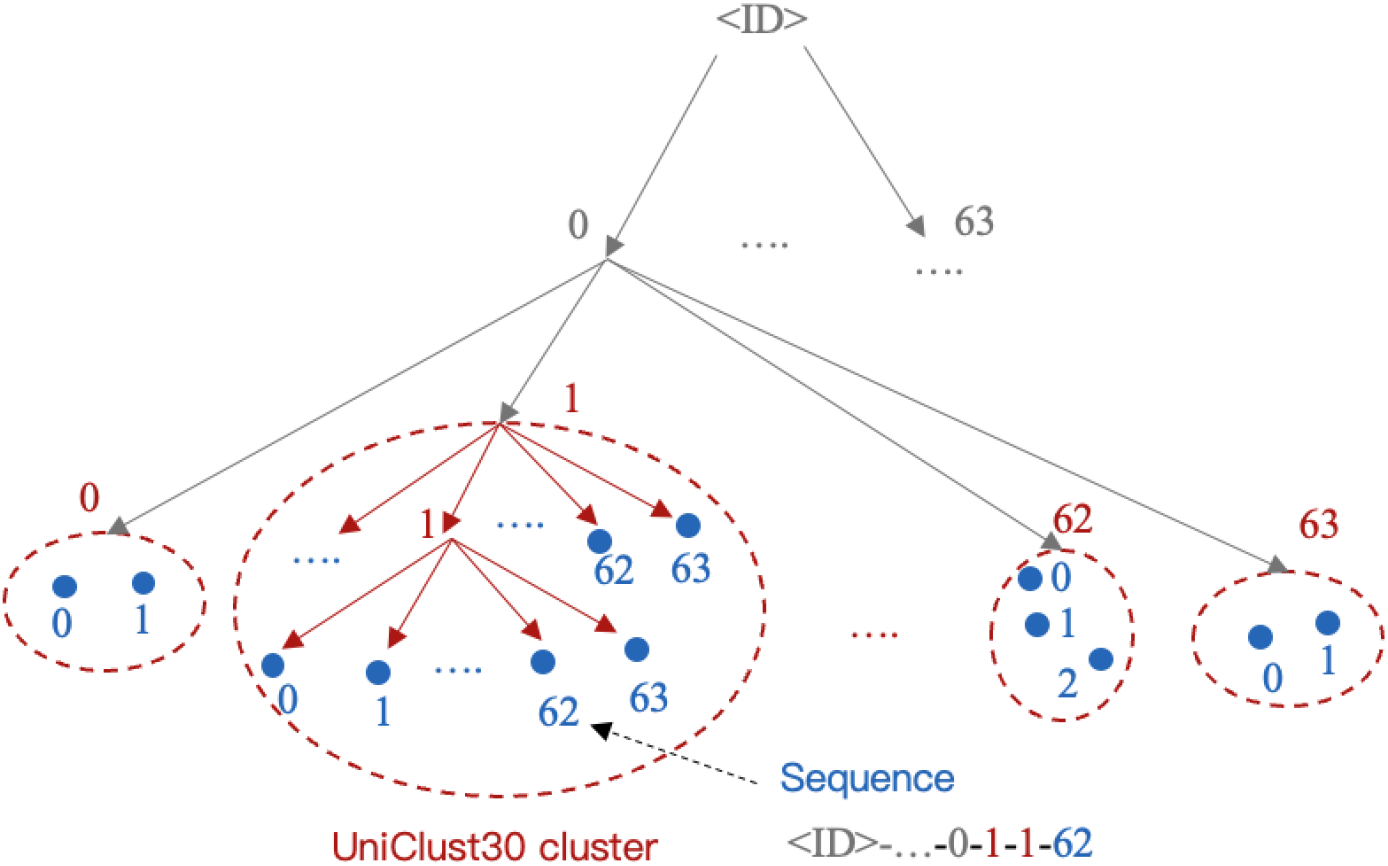
Schematic Diagram of Hierarchical ID of UniClust30 Sequences. The UC30 sequences are organized into a tree structure with a branching factor of 64. Each leaf node represents an individual sequence, while each UC30 cluster corresponds to an internal node of the tree. The hierarchical ID of a sequence is determined by traversing from the root node to the corresponding leaf node.

### 2.2. AIDO.RAGPLM

We fine-tuned a pretrained masked language model with 3-billion parameters (MLM-3B, (Cheng et al., 2024)) using MSA data by concatenating the query sequence with homologous sequences (see Figure 1). We introduced several modifications to the standard BERT masking strategy (Devlin et al., 2019): (1) We randomly sampled 0.05 *× L* span positions from a query sequence of length *L*, with span lengths following a geometric distribution (*p*=0.2), and capped the maximum length at 10. Our experiments revealed that this settings lead to an average of 15% of the query tokens were masked. (2) To prevent information leakage, when a residue was selected, all residues at the same index across all sequences (the column of the MSA matrix) were also masked. (3) When a column of MSA was selected for masking, the entire column was replaced with the <MASK> token in 80% of cases, with random amino acids in 10% of cases, and remained unchanged in the remaining 10% of cases. To help the model distinguish which tokens are from the same chain and which tokens have the same residue index, we use 2D rotary position embedding (Chen et al., 2024a; Su et al., 2023) to encode the tokens (see Figure 4 and Appendix E). For the details of training parameters, please refer to Table 6.

**Figure 4.**
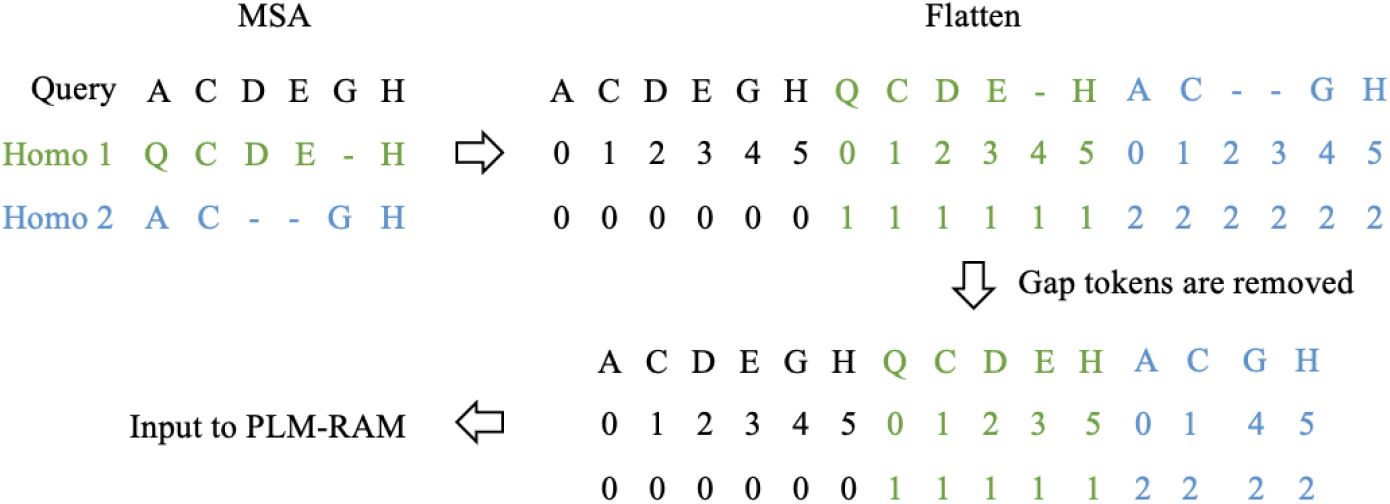
Schematic Diagram of AIDO.RAGPLM input.

### 2.3. AIDO.RAGFold

Inspired by ESMFold (Lin et al., 2023), we use AIDO.RAGPLM as a feature extractor, and added the folding trunks (AlphaFold2 Evorformer without the column attention module) and Structure modules as a head to enable end-to-end protein structure prediction. During training, we also fine-tuned the AIDO.RAGPLM base model using LoRA (Rank=16, Alpha=16). We experimented with various numbers of folding trunks and found that 24 blocks were enough, which is half the number used in AlphaFold2 and ESMFold. Additionally, we replaced the ReLU activation function with GEGLU (Shazeer, 2020) in the transition module to enhance model performance. Our training procedure consists of two phases: initial training and fine-tuning. Detailed training parameters are provided in Appendix 7. Please refer to Appendix E for the details of the data description, model training and inference.

## 3. Results

Please refer to Appendix F for details of test datasets.

### 3.1. Comparing MSA retriever and HHblits

We employed HHblits and our MSA retriever to obtain MSAs of the test sequences from the UC30 database. For MSA retriever, we experimented with two sets of parameters: (1) beam search to generate 20 UC30 clusters; (2) Top-K (K=10) sampling for 64 UC30 clusters. The results were combined and used as input for AlphaFold2 (checkpoint: `model_3_ptm`). Table 5 demonstrates that although our results are not as favorable as those obtained with HHblits in terms of TM-score, our method is approximately 45 to 85 times faster. To address the issue of missing targets in the retriever (Depth *≤* 10), we combined the MSA retriever with HHblits. For samples with a depth of less than 10 in the retriever’s results, we used HHblits to retrieve the MSA again. We found that the TM-score is comparable across four datasets when using HHblits, while still maintaining a 5 to 70-fold increase in speed.

### 3.2. AIDO.RAGPLM

#### Perplexity (PPL)

We randomly replace 15% of the tokens in the sequence with `<MASK>` token. For MSA sequences, residues at the same index (the column of MSA) of masked query are also masked. We then calculate the perplexity of the masked tokens from the query sequence using ESM2-3B, MLM-3B, and AIDO.RAGPLM models. Table 4 shows that PLMRAG has the lowest PPL across all datasets, and as the number of homologous sequences increases, the PPL decreases further.

#### Unsupervised Contact Prediction

Following the methodology of (Rao et al., 2021), we randomly selected 20 chains as the training set and obtained *H × L* attention maps from the model, where *H* is the number of heads and *L* is the number of layers. Each attention map was symmetrized and adjusted using the Average Product Correction (APC) independently. Residue pairs with a distance of less than 8A° were defined as contacts. Logistic regression was employed to predict whether a residue pair is a contact (distance less than 8A°) using the *H × L* features as input. For AIDO.RAGPLM, only the attention map of the query sequence part was utilized. As shown in Table 1, AIDO.RAGPLM outperforms the two single-sequence models, despite its base model, MLM-3B, performing worse than ESM-3B on the CAMEO and Recent datasets.

**Table 1.** Unsupervised contact prediction.

|  |  | ESM-3B | MLM-3B | RAGPLM |
| --- | --- | --- | --- | --- |
| Top L | CASP14 | 0.357 | 0.350 | **0.389** |
|  | CASP15 | 0.420 | 0.427 | **0.451** |
|  | CAMEO | 0.493 | 0.483 | **0.513** |
|  | Recent | 0.452 | 0.433 | **0.477** |
| Top L/5 | CASP14 | 0.348 | 0.365 | **0.396** |
|  | CASP15 | 0.381 | 0.380 | **0.415** |
|  | CAMEO | 0.444 | 0.441 | **0.470** |
|  | Recent | 0.436 | 0.403 | **0.443** |

#### Supervised Contact Prediction

We utilized the contact prediction dataset from trRosetta (Yang et al., 2020) to fine-tune the model. For all models, qkvo LoRA (Hu et al., 2021) and MLP LoRA were applied with (Rank=16, Alpha=16). The batch size was set to 8, and training was conducted for 25,000 steps. The checkpoint with the highest validation Top *L/*5 accuracy was used to evaluate the model. As shown in Table 2, AIDO.RAGPLM outperforms ESM2-3B and MLM-3B on both the validation and test sets.

**Table 2.** Result of supervised contact prediction and fitness prediction. Supervised contact prediction: 1,512 samples for validation set and test set. Fitness prediction: Spearman correlation coefficients of Deep Mutational Scanning (DMS) assays from ProteinGym. The column labeled “All” includes sequences with single and multiple mutations, while the column labeled “Single” includes sequences with only a single mutation. The data size is 207.

|  | Contact |  | Fitness |  |
| --- | --- | --- | --- | --- |
|  | Validation | Test | All | Single |
| ESM2-3B | 0.931 | 0.915 | 0.439 | 0.426 |
| MLM-3B | 0.916 | 0.910 | 0.430 | 0.408 |
| PLMRAG | **0.938** | **0.927** | **0.462** | **0.437** |

#### ProteinGym zero-shot prediction

We obtained the substitutions dataset of Deep Mutational Scanning (DMS) assays from the ProteinGym website (Notin et al., 2023). For each mutation *t*_*wt*_ *→ t*_*mut*_, we replace the wildtype token *t*_*wt*_ with a special `<MASK>` token. We then computed the log ratio *− log*(*P*_*θ*_(*t*_*mut*_)) − *log*(*P*_*θ*_(*t*_*wt*_)), where *P*_*θ*_(*t*_*mut*_) represents the model’s probability of the mutated token given the other tokens as input. To evaluate the model’s performance, we calculated the Spearman correlation coefficient between the log ratio and the “DMS score” from the downloaded tables. As shown in Table 2, the AIDO.PLMRAG model achieved a higher score compared to the other two single-sequence PLMs.

### 3.3. AIDO.RAGFold

We conducted a comparative analysis of TM-scores and runtime between AIDO.RAGFold and AlphaFold2 (checkpoint: model 3 ptm) using HHblits retrieved MSAs as input. The number of recycle (*N*_*recycle*_) was fixed at three, and the maximum context length for RAG was constrained up to 25,600. Both AlphaFold2 and AIDO.RAGFold were executed with varying *N*_*ensemble*_ (1, 2 and 4), resulting in AIDO.RAGPLM processing 4, 8, and 16 different MSAs, respectively (see Algorithm 1). Table 3 and Table 12 presents the TM-score of the two models. Table 10 presents the inference time, RMSD and LDDT. Our findings indicate that:

**Table 3.** TM-scores of AlphaFold2, AIDO.RAGFold, and ESM-Fold on four test datasets. HHblits MSAs were used as input for AlphaFold2 and AIDO.RAGFold. *N*_*ensemble*_ = 4.

| Dataset | AF2 | AIDO.RAGFold | ESMFold |
| --- | --- | --- | --- |
| CASP14 | 0.767 | **0.776** | 0.696 |
| CASP15 | **0.728** | 0.726 | 0.639 |
| CAMEO | 0.864 | **0.871** | 0.854 |
| Recent | **0.824** | 0.823 | 0.775 |

#### MSA ensembling enhances AIDO.RAGFold’s performance

This improvement is primarily due to RAG’s limited MSA context usage. Increasing *N*_*ensemble*_ allows AIDO.RAGFold to use more homologous sequence information.

#### AIDO.RAGFold’s performance is comparable to AlphaFold2

AIDO.RAGFold demonstrates a significantly faster inference speed, ranging from 8 times faster.

#### AIDO.RAGFold outperforms ESMFold

The inclusion of MSA significantly boosts AIDO.RAGFold’s performance compared to ESMFold.

To investigate the impact of the number of MSAs on AIDO.RAGFold’s structural prediction accuracy, we randomly sampled 0, 5, 10, 25, 50, and 100 sequences from the HHblits MSA as input for both AlphaFold2 and AIDO.RAGFold. Table 11 and Figure 2 illustrate that AIDO.RAGFold’s TM-scores surpass those of AlphaFold2 when the number of MSAs is limited. For instance, using the Recent PDB dataset, AIDO.RAGFold outperforms AlphaFold2 by margins of 0.420, 0.155, 0.070, 0.016, and 0.007 for 0, 5, 10, 25, 50, and 100 MSAs, respectively. However, it is noteworthy that without any MSA input, AIDO.RAGFold’s performance lags behind ESM-Fold. Nevertheless, providing more than 5 MSAs enables AIDO.RAGFold to match ESMFold’s performance, with the exception of the CAMEO dataset.

## 4. Conclusion

Our study introduces a novel MSA retrieval method based on ID generation, significantly accelerating MSA acquisition compared to traditional approaches. Utilizing this method, we expanded the existing MSA dataset and trained an MSA retrieval-enhanced protein language model. Our findings indicate that this model outperforms single-sequence models in tasks such as contact prediction and fitness prediction. Furthermore, we employed the embeddings from this language model for downstream end-to-end structure prediction, achieving results comparable to AF2, but with an approximately eightfold increase in speed. Notably, in scenarios with insufficient MSAs, our model substantially surpasses AF2, underscoring the critical importance of pre-trained models.

## A. Retrieval Augmented Protein Language Models

Recent advancements in protein language models have attempted to integrate multiple homologous sequences for training. For example, the MSA Transformer (Rao et al., 2021), a model with 150 million parameters, utilizes aligned homologous sequences as input and employs self-supervised learning through random masking. This model has demonstrated superior performance compared to single-sequence models in downstream tasks such as fitness and contact prediction. Similarly, PoET (au2 & Bepler, 2023), an autoregressive generative model, concatenates unaligned sequences and trains them using next-token prediction. This enables the generation of entirely new sequences within the same family and the prediction of variant fitness. RSA (Ma et al., 2023) retrieves homologous sequences of the query using its dense sequence retriever and aggregates the information from the query and homologous sequences in pairs for downstream task prediction. This method not only achieves a retrieval speed significantly faster than traditional MSA methods but also delivers superior results in tasks such as fold classification, contact prediction, and localization. ProtMamba (Sgarbossa et al., 2024), leveraging the Mamba framework, extends the maximum sequence length up to 131k. By integrating autoregressive modeling and masked language modeling (MLM) with a fill-in-the-middle objective, ProtMamba can generate protein sequences and be utilized for downstream tasks such as fitness prediction.

## B. MLM-3B model and CLM-3B model

The MLM-3B model (Cheng et al., 2024)) is a transformer encoder framework with 2.8 billion parameters. We utilize the same hyperparameters as ESM-3B, specifically: 36 layers, 40 heads, a hidden size of 2560, and an FFN hidden size of 6832. The training data is a mixture of the UniRef database and ColabFoldDB. We follow the BERT masking strategy: 15% of the tokens are selected for masking, with 80% replaced by special MASK tokens, 10% replaced by random amino acids, and the remaining 10% left unchanged. The learning rate schedule includes a 3% warm-up phase from 0 to 2.5e-4 followed by cosine decay from 2.5e-4 to 2.5e-5. Please refer to Table 6 for detailed information about MLM-3B. We train on 1,000 billion tokens and evaluate the model on two out-of-distribution datasets, with maximum identity to the training set being less than 0.9 and 0.5, respectively. The results are presented in Table 5.

The CLM-3B model (Cheng et al., 2024)) is a transformer decoder framework. It shares the same hyperparameters as the MLM-3B model and is trained on the same dataset. Our approach follows the training methodology of GPT, predicting the next token based on the given prefix. For detailed information about CLM-3B, please refer to Table 6.

## C. MSA retriever

As described in the main text, training the MSA retriever involves three steps. Below, we detail the methods for each step.

### C.1. Construct Hierarchical IDs for Each Sequence in UniClust30 via Hierarchical K-means Clustering of Embedding

The UC30 database (v2021_03) comprises 29 million clusters containing a total of 263 million sequences. The hierarchical ID is a multi-layer tree structure (see Figure 3), with each node having no more than 64 child nodes. Each leaf node corresponds to a sequence in UC30. To ensure the hierarchical ID reflects sequence similarity semantics (e.g., the similarity between 23-43-52-0 and 23-43-52-1 is higher than that between 23-43-52-0 and 23-43-34-5), we assign the ID tokens by clustering the embedding of sequences. So we use the MLM-3B model to generate embedding (dimension of 2560) for the 263 million sequences.

The ID of a sequence consists of two parts: (1) ID_center: derived from clustering the 29 million cluster centers; (2) ID_member: derived from clustering the members within a UC30 cluster.

#### ID_center

For each UC30 cluster, the longest sequence is selected as the representative sequence, and its embedding is used as the representative embedding of the cluster. We perform hierarchical K-means clustering (K=64) on the 29 million representative embedding, resulting in a tree with a degree of 64. We label all child nodes of each node from 0 to 63. Thus, for any node, we traverse from the root node to it in sequence to obtain its hierarchical ID, which is ID_center.

#### ID_member

For UC30 clusters with more than 64 members, we perform the same hierarchical K-means clustering on all members’ embeddings to build ID_member.

The final ID for each sequence is obtained by concatenating ID_center and ID_member.

### C.2. Fine-tune the CLM-3B Model to Memorize the ID of Each Sequence

We first build a Seq-ID dataset from UC30 dataset. Each sample comprises the query sequence, a special `<ID>` token, the hierachical ID tokens, and an `<EOS>` token. The CLM-3B model is trained with 500 billion tokens. After training, the model can generate the ID tokens with the query sequence and the `<ID>` token as a prefix until the `<EOS>` token or the UC30 cluster level token (purple circle in Figure 3). The learning rate is warmed up from 0 to 2.0e-5 for the first 2.5% of training tokens and then decays to 0 using a cosine schedule.

### C.3. Continue to fine-tune the model to Generalize to IDs of Homologous Sequences

We use HHblits to search for MSAs from UC30 using UniRef50 (UR50) as query sequences, obtaining 23.7 million MSAs. We refer to this dataset as HHblits_MSA. When fine-tuning CLM-3B on this dataset, each sample comprises a query sequence, a special `<ID>` token, the ID tokens (randomly sampled from its homologous sequences), and a `<EOS>` token. We train 10 billion tokens on this dataset. Please refer to Table 6 for detailed information.

## D. AIDO.RAGPLM training dataset

We utilized sequences from UniRef50 as queries to search for homologous sequences in UniClust30, subsequently constructing multiple sequence alignments (MSAs). UniRef50 comprises a total of 53.6 million sequences. Using HHblits, we searched all sequences, identifying over 25 homologous sequences for 23.7 million of them. This dataset was directly used as the training set, referred to as HHblits MSA. The remaining 29.9 million sequences were input into MSA Retriever, resulting in 7.7 million sequences with more than 25 homologous sequences. This dataset was designated as Retriever MSA. During training, AIDO.RAGPLM randomly sampled from the two datasets with probabilities of 0.75 and 0.25, respectively. Detailed information is provided in Figure 8.

## E. Detailed description of AIDO.RAGFold architecture and inference

We used the PDB database (release prior to January 1, 2024), the AlphaFold Database (with mean pLDDT *≥* 90) and OpenProteinSet residues with pLDDT *≥* 90 (Ahdritz et al., 2023) as the training set. Detailed information about the data is provided in Table 9. We ensured that all samples with sequence identity greater than 0.5 with the test set were excluded. The open-source OpenFold framework was employed to train our RAG-Fold model.

To feed the query tokens (∈ ℝ^*L*^, where *L* is the length of the query sequence) and MSA tokens (∈ ℝ^*N ×L*^, where *L* is the length of the query sequence) into the AIDO.RAGPLM model, the MSA tokens are flattened into the shape of ℝ^*NL*^. We initialize a 2D positional encoding (∈ ℝ^2*×L*^), where the first dimension represents the residue index for each sequence and the second dimension represents the sequence index (Chen et al., 2024a). To reduce the length of the sample, we remove *G* gap tokens that contain no information in the sequence. This adjustment changes the dimension of the sample to ℝ^*NL−G*^ and the dimension of the positional encoding to ℝ^2*×*(*NL−G*)^. Figure 4 illustrates this process.

The output of the AIDO.RAGPLM model includes the embeddings of homologous sequences. We retain only the hidden states corresponding to the query tokens and input them into the downstream modules. Linear modules are employed to transform these hidden states into the MSA representation and Pair representation of folding trunks. For a detailed description, please refer to Algorithm 1.

During inference, due to the limitation of the input sample length (up to 25,600), the information from homologous sequences that the AIDO.RAGPLM model can utilize is restricted. To address this, we adopted the MSA ensembling method from AlphaFold2. Specifically, we sample a subset of up to 25,600 sequences from the all MSA sequences each time and run the AIDO.RAGPLM *N*_*ensemble*_ times to average the resulting representations. This approach enables us to maximize the utilization of information from homologous sequences.

## F. Test datasets

- **CASP14** (N=50): Protein targets obtained from the CASP14 website, accompanied by ground-truth structures.
- **CASP15** (N=53): Protein targets sourced from the CASP15 website, with corresponding ground-truth structures.
- **CAMEO** (N=194): Protein domains retrieved from the CAMEO website, covering the period from July 1, 2021, to June 1, 2022.
- **Recent PDB** (N=107): Protein chains extracted from the PDB database, with release dates ranging from January 1, 2024, to July 1, 2024. The following criteria were applied to filter the chains: (1) a length range between 50 and 1500 residues; (2) exclusion of sequences containing non-standard amino acid types; (3) removal of sequences with repeat fragments, defined as having a bi-gram entropy greater than 4; (4) exclusion of sequences with more than 50% identity to the training set; (5) clustering of sequences at a 50% identity cutoff, selecting one representative sequence per cluster.

**Table 4.** Perplexity of various models and inputs across six sequence datasets. (N) denotes the dataset size, while (D) represents the number of homologous sequences used as input for AIDO.RAGPLM.

|  | CASP14 | CASP15 | CAMEO | Recent PDB | MaxID0.5 | MaxID0.9 |
| --- | --- | --- | --- | --- | --- | --- |
| N | 50 | 53 | 194 | 107 | 5,012 | 6,907 |
| ESM2-3B | 10.658 | 5.963 | 5.959 | 6.223 | 10.753 | 6.703 |
| MLM-3B | 8.905 | 6.355 | 5.631 | 5.671 | 10.959 | 6.816 |
| RAGPLM (D=1) | 10.114 | 6.599 | 6.321 | 6.357 | 10.718 | 6.816 |
| RAGPLM (D=8) | 9.303 | 6.360 | 6.212 | 5.999 | 10.222 | 6.494 |
| RAGPLM (D=16) | 8.980 | 6.167 | 6.167 | 5.666 | 9.989 | 6.317 |
| RAGPLM (D=64) | 8.724 | 5.803 | 5.995 | 5.329 | 9.381 | 5.840 |
| RAGPLM (D=128) | 8.391 | **5.612** | 5.707 | 5.296 | **9.341** | **5.833** |
| RAGPLM (D=256) | **8.072** | 5.741 | **5.635** | **5.266** | 9.359 | 5.871 |

**Table 5.** Performance Comparison of Various MSA Search Tools. In the case of Ours + hhblits, Ours MSAs with a depth of fewer than 10 were replaced with HHblits MSAs.

| | | Average time (s) | #(Depth $\geq 100$) ↑ | #(Depth $\leq 10$) ↓ | AlphaFold2<br>TM-score ↑ |
| --- | --- | --- | --- | --- | --- |
| CASP14 | hhblits | 899 | 27 | 9 | **0.757** |
|  | Ours | 19 | 31 | 11 | 0.696 |
|  | Ours + hhblits | 172 | 31 | 7 | 0.748 |
| CASP15 | hhblits | 2928 | 47 | 2 | **0.731** |
|  | Ours | 32 | 42 | 8 | 0.668 |
|  | Ours + hhblits | 94 | 46 | 2 | 0.723 |
| CAMEO | hhblits | 2761 | 172 | 5 | **0.865** |
|  | Ours | 43 | 163 | 24 | 0.831 |
|  | Ours + hhblits | 127 | 171 | 5 | 0.862 |
| Recent PDB | hhblits | 3138 | 91 | 1 | 0.824 |
|  | Ours | 43 | 94 | 10 | 0.783 |
|  | Ours + hhblits | 104 | 96 | 1 | **0.827** |

**Table 6.** Detailed training information of MLM-3B, CLM-3B, MSA Retriever and AIDO.RAGPLM.

|  | MLM-3B | CLM-3B | Retriever<br>Step 1 | Retriever<br>Step 2 | RAGPLM |
| --- | --- | --- | --- | --- | --- |
| Training data | UniRef +<br>ColabFoldDB | UniRef +<br>ColabFoldDB | UniClust30 | HHblits_MSA | HHblits_MSA<br>Retriever_MSA |
| Initial params | Random | Random | CLM-3B | Retriever<br>Step 1 | MLM-3B |
| Learning rate | 2.5e-4 | 1.2e-4 | 2e-4 | 1.2e-4 | 1e-4 |
| Training tokens | 1000B | 2300B | 300B | 10B | 100B |
| Batch size | 2560 | 2048 | 2048 | 1024 | 256 |
| Micro batch size | 4 | 4 | 4 | 4 | 1 |
| Sample length | 1024 | 2048 | 2048 | 1024 | 12800 |
| Attention | Bi-directional | Causal | Causal | Causal | Bi-directional |

**Table 7.** Detailed training information of AIDO.RAGFold.

|  | Initial training | Fine-tuning |
| --- | --- | --- |
| Sequence crop size | 256 | 368 |
| Maximum context length of RAG | 16,384 | 12,800 |
| Exponential moving average | Enabled | Enabled |
| Learning rate of LoRA | A: 1e-4, B: 1.6e-3 | A: 1e-4, B: 1.6e-3 |
| Learning rate of folding trunks Structural modules | First 90%: 1e-3<br>Last 10%: 5e-4 | 5e-4 |
| Batch size | First 90%: 128<br>Last 10%: 256 | 256 |
| Warm up | First 2000 steps | N/A |
| Structural violation loss weight | 0 | 0.1 |
| ”Experimentally resolved” loss weight | 0 | 0.01 |
| Training samples (million) | 10 | 6 |

**Table 8.** Training data of AIDO.RAGPLM. 23.7 million MSAs are collected by HHblits and 7.7 million MSAs are collected by MSA Retriever.

|  | #Seqs | #Query tokens | Sample Weight |
| --- | --- | --- | --- |
| HHblits_MSA | 23.7M | 6.5B | 0.75 |
| Retriever_MSA | 7.7M | 2.4B | 0.25 |

**Table 9.** Training data of AIDO.RAGFold.

| Dataset | #Chains | #Clusters | Sample ratio |
| --- | --- | --- | --- |
| PDB | 440,952 | 34,961 | 25% |
| AlphaFold DB Distil | 4,457,794 | 1,829,120 | N/A |
| OpenProteinSet Distil | 259,343 | 242,079 | N/A |
| Distil mixed | 4,711,621 | 2,002,005 | 75% |

**Table 10.**
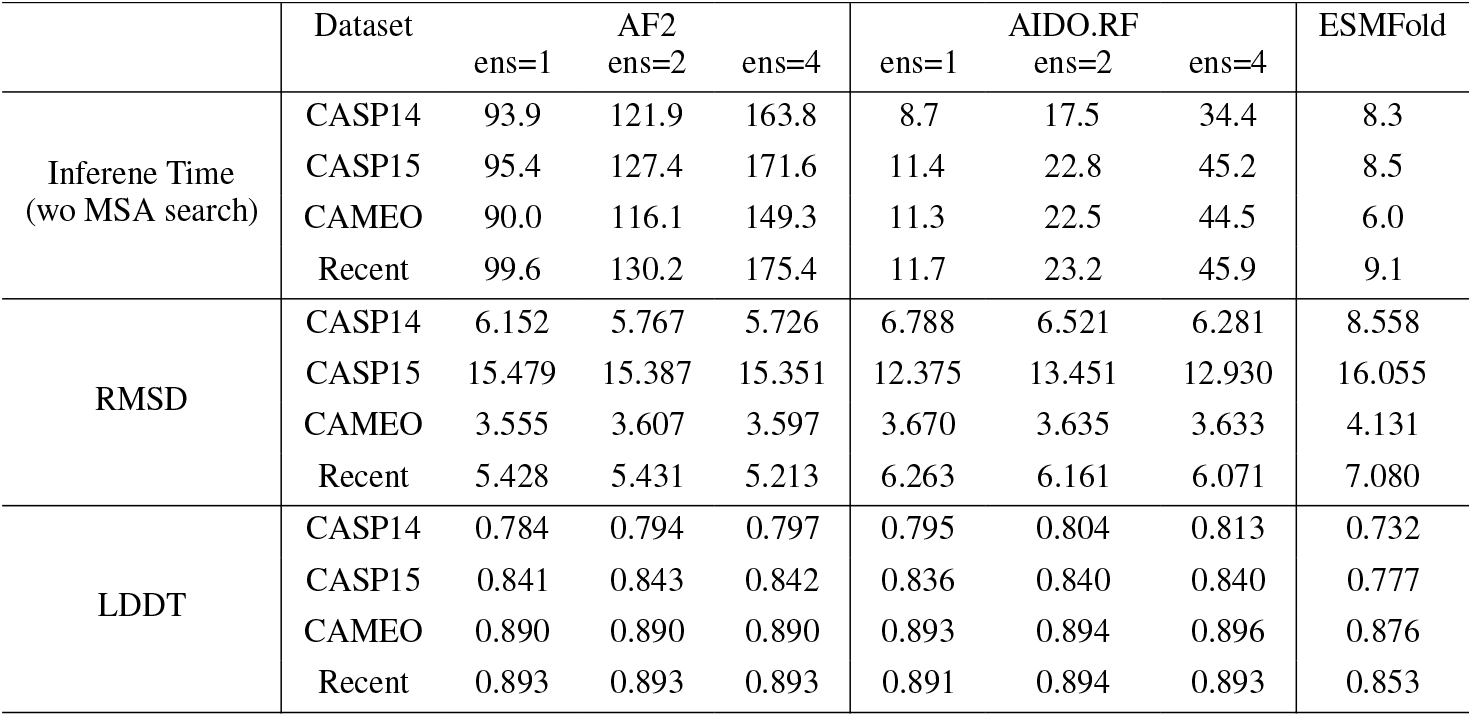
Inference time, RMSD and LDDT of AlphaFold2 (AF2), AIDO.RAGFold (AIDO.RF), and ESMFold on four test datasets.

**Table 11.** TM-scores of AlphaFold2, AIDO.RAGFold, and ESMFold on four test datasets with limited MSA sequences as input.

|  | #MSA=0 |  | #MSA=5 |  | #MSA=10 |  | #MSA=25 |  | #MSA=50 |  | #MSA=100 |  | ESMFold |
| --- | --- | --- | --- | --- | --- | --- | --- | --- | --- | --- | --- | --- | --- |
|  | AF2 | RAG | AF2 | RAG | AF2 | RAG | AF2 | RAG | AF2 | RAG | AF2 | RAG |  |
| CASP14 | 0.298 | 0.604 | 0.584 | 0.692 | 0.672 | 0.728 | 0.716 | 0.735 | 0.726 | 0.744 | 0.740 | 0.748 | 0.696 |
| CASP15 | 0.290 | 0.624 | 0.583 | 0.652 | 0.614 | 0.666 | 0.645 | 0.678 | 0.666 | 0.697 | 0.701 | 0.697 | 0.639 |
| CAMEO | 0.330 | 0.787 | 0.690 | 0.822 | 0.777 | 0.834 | 0.820 | 0.844 | 0.840 | 0.851 | 0.843 | 0.853 | 0.854 |
| Recent | 0.292 | 0.712 | 0.618 | 0.773 | 0.714 | 0.784 | 0.769 | 0.798 | 0.790 | 0.806 | 0.800 | 0.807 | 0.775 |

**Table 12.** TM-scores of AlphaFold2, AIDO.RAGFold, and ESMFold on four test datasets. HHblits MSAs were used as input for AlphaFold2 and AIDO.RAGFold. “ens” denotes the number of MSA ensembles.

| Dataset | AF2 |  |  | AIDO.RAGFold |  |  | ESMFold |
| --- | --- | --- | --- | --- | --- | --- | --- |
|  | ens=1 | ens=2 | ens=4 | ens=1 | ens=2 | ens=4 |  |
| CASP14 | 0.754 | 0.766 | 0.767 | 0.752 | 0.764 | **0.776** | 0.696 |
| CASP15 | 0.725 | 0.727 | **0.728** | 0.722 | 0.727 | 0.726 | 0.639 |
| CAMEO | 0.864 | 0.863 | 0.864 | 0.868 | 0.869 | **0.871** | 0.854 |
| Recent | **0.824** | 0.823 | **0.824** | 0.820 | 0.823 | 0.823 | 0.775 |

### Algorithm 1 AIDO.RAGFold

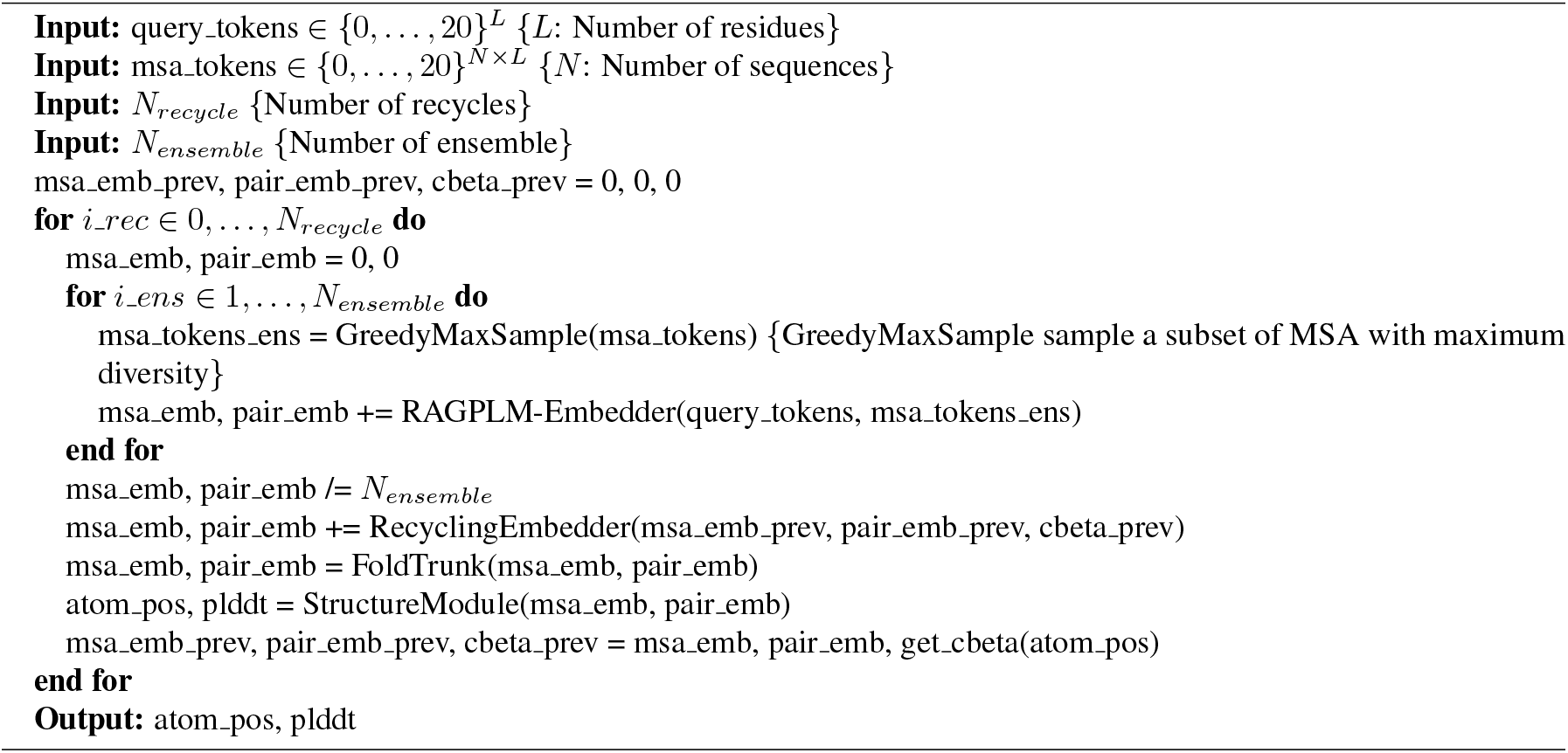

### Algorithm 2 RAGPLM-Embedder

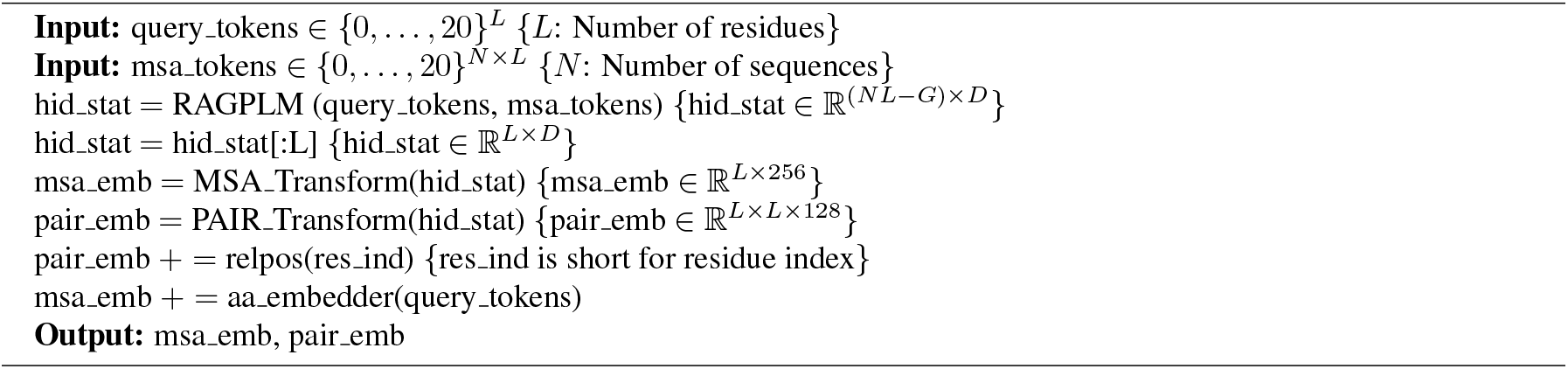

### Algorithm 3 MSA_Transform

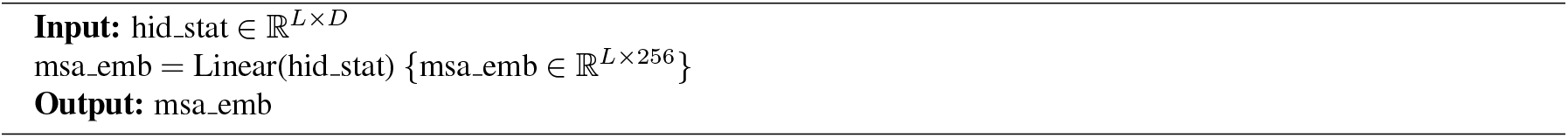

### Algorithm 4 PAIR_Transform

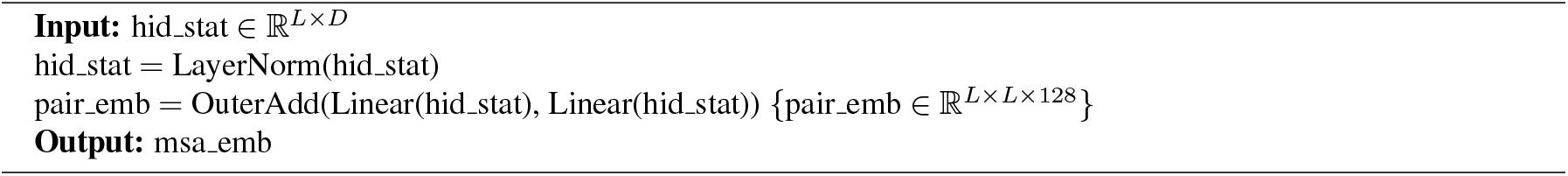

